# Natural image statistics at depth edges modulate perceptual stability

**DOI:** 10.1101/2020.04.05.026724

**Authors:** Zeynep Başgöze, David N. White, Johannes Burge, Emily A. Cooper

## Abstract

Binocular fusion relies on matching points in the two eyes that correspond to the same physical feature in the world. However, not all world features are binocularly visible. In particular, at depth edges parts of a scene are often visible to only one eye (so-called *half occlusions*). Accurate detection of these monocularly visible regions is likely to be important for stable visual perception. If monocular regions are not detected as such, the visual system may attempt to binocularly fuse non-corresponding points, which can result in unstable percepts. We investigated the hypothesis that the visual system capitalizes upon statistical regularities associated with depth edges in natural scenes to aid binocular fusion and facilitate perceptual stability. By sampling from a large set of stereoscopic natural image patches, we found evidence that monocularly visible regions near depth edges in natural scenes tend to have features more visually similar to the adjacent binocularly visible background region than to the adjacent binocularly visible foreground. The generality of these results was supported by a parametric study of three-dimensional (3D) viewing geometry in simulated environments. In two perceptual experiments, we examined if this statistical regularity may be leveraged by the visual system. The results show that perception tended to be more stable when the visual properties of the depth edge were statistically more likely. Exploiting regularities in natural environments may allow the visual system to facilitate fusion and perceptual stability of natural scenes when both binocular and monocular regions are visible.

**Precis:** We report an analysis of natural scenes and two perceptual studies aimed at understanding how the visual statistics of depth edges impact perceptual stability. Our results suggest that the visual system exploits natural scene regularities to aid binocular fusion and facilitate perceptual stability.

## Introduction

In animals with binocular vision, the two eyes’ images must be fused to obtain a single percept of the world. Because the eyes capture slightly different views, fusion requires determining which retinal points in the two eyes correspond to the same physical feature in the world (**Figure 1**, connected yellow squares). Due to ambiguities in this retinal correspondence, binocular fusion is computationally demanding. When the visual system fails to fuse the two eyes’ images, observers can experience *unstable* percepts. This experience is often called binocular rivalry, because percepts can alternate between the content visible to each of the two eyes (Levelt, 1965). To reduce the computational complexity of binocular fusion and obtain a stable percept of the world, the visual system is known to exploit statistical regularities that are present in binocular images of natural scenes (Burge, Fowlkes, & Banks, 2010; Burge & Geisler, 2014; Cooper, Burge, & Banks, 2011; Cooper & Norcia, 2015; Gibaldi, Canessa, & Sabatini, 2017; Goncalves & Welchman, 2017; Hibbard & Bouzit, 2005; Samonds, Potetz, & Lee, 2012; Sprague, Cooper, Tošić, & Banks, 2015).

**Figure 1.**
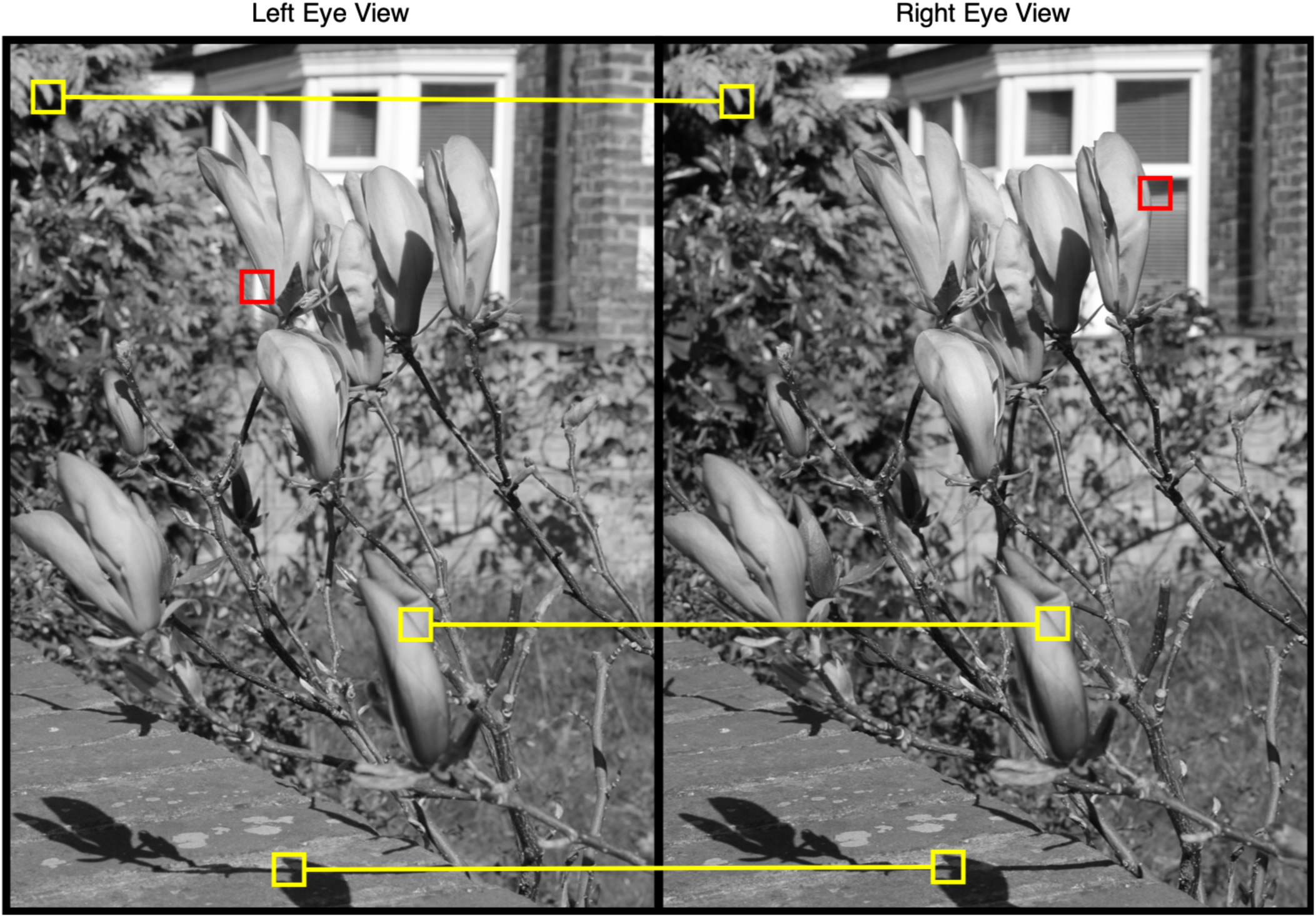
Binocular and monocular retinal projections during natural vision. A stereo view of a natural scene is shown, representing the images cast on the left and right retinas of an observer. To obtain a single percept of the world, these two views must be fused. Examples of binocularly corresponding features are highlighted with pairs of yellow squares connected by lines. Note that these features have variable binocular offsets and many similar features are visible to the two eyes, making fusion computationally challenging. Isolated red squares depict monocularly visible regions associated with depth edges. (Image courtesy of Deviant Art user aegiandyad, Creative Commons Attribution 3.0 License).

Not all features in the world, however, create binocularly matched retinal projections. Near depth edges, parts of a scene are often visible to only one eye (**Figure 1**, isolated red squares). If these monocular regions are not recognized, the visual system may attempt to fuse regions that do not binocularly correspond, resulting in unstable percepts (Forte, Peirce, & Lennie, 2002; Hoffman & Banks, 2010; Shimojo & Nakayama, 1990). The visual system should therefore exploit statistical regularities in natural images that facilitate detection of monocular features. It is likely that strong statistical regularities are associated with monocular regions, because of the geometric constraints imposed by depth edges and the visual properties of the surfaces on either side of the depth edge (e.g., Anderson, 1994; Gillam & Borsting, 1988; Grove & Ono, 1999; Nakayama & Shimojo, 1990; Tyler, Christopher & Kontsevich, 1995; see Harris & Wilcox, 2009 for an extensive review). Interestingly, although the statistical properties of binocularly visible content in natural scenes are under active study (Burge & Geisler, 2014; Chauhan, Masquelier, Montlibert, & Cottereau, 2018; Goncalves & Welchman, 2017; Hibbard, 2008; Iyer & Burge, 2018; Sprague et al., 2015), the statistical properties that characterize monocularly visible regions in natural scenes have received scant attention. The current manuscript addresses this gap.

Consider three illustrative geometric configurations that can cause monocular regions in 3D scenes. The first configuration is a “background occlusion”, in which a foreground surface partially occludes an extended background surface, so that one portion of the background surface is visible to only one eye and the other portion is visible to both eyes. In this case, the appearance of the monocular region should be similar to the binocular background (**Figure 2A**). The second configuration is a “self-occlusion”, in which a foreground surface occludes some portion of itself, rendering some portion of the foreground surface visible to only one eye. In this case, the appearance of the monocular region should be similar to the binocular foreground (**Figure 2B**). The third configuration is a “hidden surface”, in which a foreground surface occludes a surface behind it such that the hidden surface is imaged by only one eye (**Figure 2C**). In this last case, the monocularly visible surface in the scene is not continuous with either the adjacent binocular foreground or the adjacent binocular background; the features in the monocular image region are thus likely to be dissimilar to those in the image regions corresponding to the binocularly visible foreground and background. Additionally, monocular regions may often be comprised of a combination of multiple surfaces (“hybrid occlusions”, see Appendix **Figure A1**). For example, when the edge of the foreground surface is curved, a single monocular region can contain some portion of the foreground directly adjacent to some portion of the background. The configurations shown in **Figure 2** thus represent extremes along a continuum of possible configurations, which represent useful starting points for geometric and statistical analyses.

**Figure 2.**
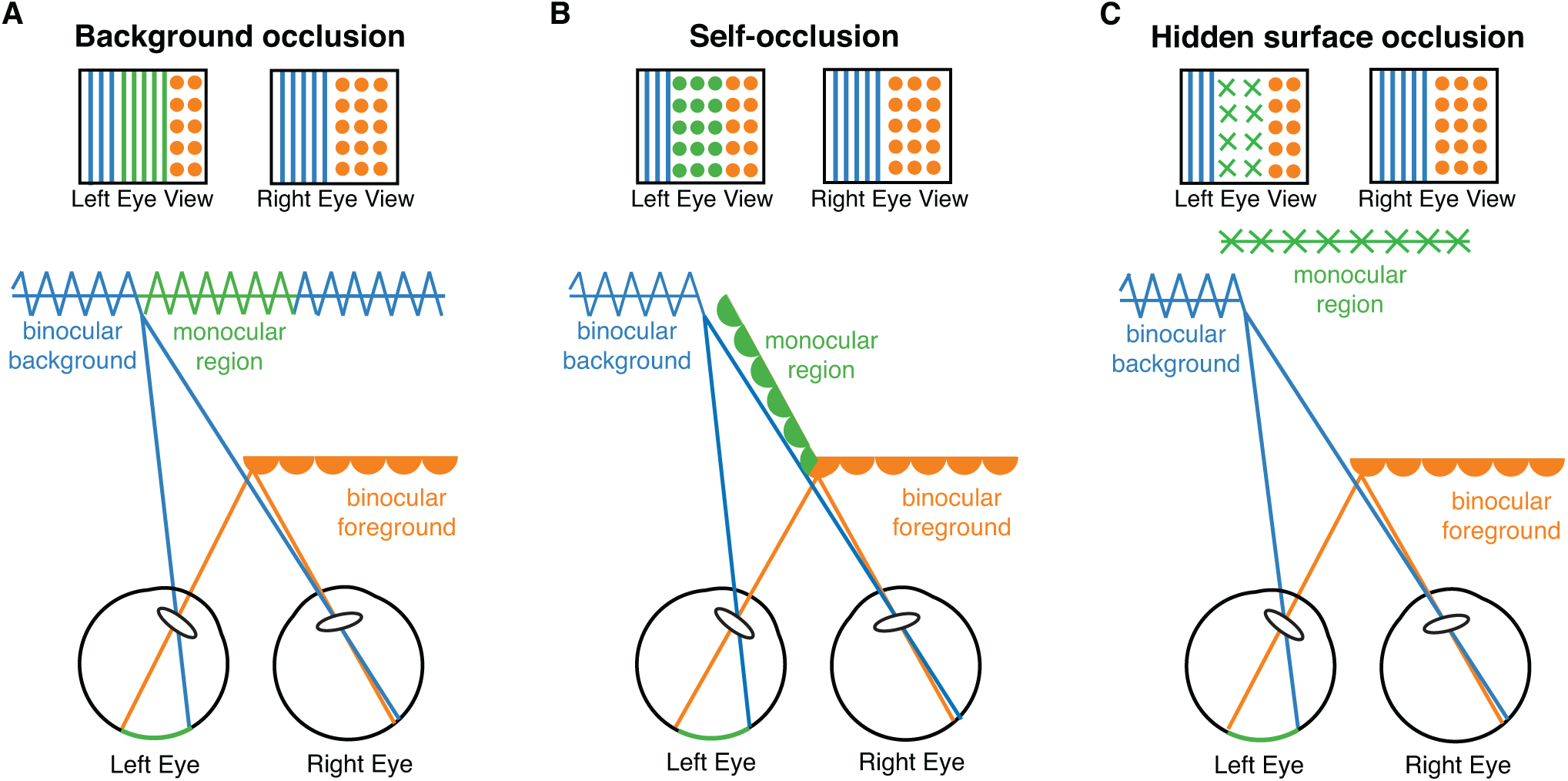
Diagrams of depth edge configurations that lead to different types of monocular regions. A local area of a scene containing a single depth edge can be subdivided into a monocularly visible region (green), the adjacent binocularly visible background (blue), and the adjacent binocularly visible foreground (orange). Patterns above each diagram illustrate the left- and right-eye views of the scene. Note that in all configurations, the right eye views are the same, and the left eye views contains a monocular region. If the depth edge is reversed, so the foreground is on the left, the monocular region will fall in the other eye. Colors are used to indicate monocular versus binocular content, not actual color differences in the scene. **A.** In a background occlusion, the left eye sees around the corner of the foreground to expose more of the background surface**. B.** In a self-occlusion, the foreground occludes itself, and the left eye sees the side of the foreground object, but the right eye does not. **C.** In a hidden surface occlusion, there is an object or surface that the foreground hides from the right eye, but the left eye can see it. Any given depth edge may contain a combination of any of these factors (see **Figure A1**).

If the visual system is adapted to process depth edge configurations that are most likely to be encountered in natural scenes, then perceptual experiments may provide clues about the corresponding statistical regularities. For example, if background occlusions are the most likely cause of monocular regions, visual systems might better process images in which the monocular region is more similar to the binocularly visible background than to the binocularly visible foreground. Consistent with this hypothesis, experiments using random dot stereograms of depth edges have shown that depth processing is faster when the texture of the monocular region matches the texture of the binocularly visible background (Gillam & Borsting, 1988; Grove & Ono, 1999). Experiments have also shown that the perceived depth of the monocular region tends to match the perceived depth of the binocularly visible background (Grove, Ben Sachtler, & Gillam, 2006; Grove, Gillam, & Ono, 2002). However, not all such studies have produced consistent results. In the only study to date (that we are aware of) that has used natural images to probe the processing and perception of monocular regions, judgments about the depth structure of the scene were found to be faster for self-occlusions than for background occlusions (Wilcox & Lakra, 2007).

In a complementary line of research, computational studies have been used to examine statistical regularities in monocular regions. For example, simulations of 3D scene geometry have shown that the probability of a point being monocularly visible varies as a function of distance in scenes composed of cluttered surfaces (Langer & Mannan, 2012; Langer, Zheng, & Rezvankhah, 2016). However, the image cues that may be available to the visual system for identifying these monocular regions in typical natural environments have not been examined. Thus, given the current literature, we do not yet fully understand the visual processing and perception of monocular regions near depth edges, particularly in natural scenes.

In recent decades, a substantial literature has developed that links the statistics of natural images and the geometry of natural scenes to the structure and function of the human visual (see (Geisler, 2008) for an extensive review). Here, we investigate the hypothesis that depth edges in natural scenes contain statistical regularities that the visual system capitalizes upon to facilitate perceptual stability. We sampled and analyzed a large set of stereoscopic image patches containing depth edges from a natural image database with co-registered distance measurements (Burge, McCann, & Geisler, 2016). In two perceptual experiments, human observers viewed the sampled patches and rated the visual stability of the depicted scene near each depth edge. The results showed that perception was relatively more stable when the properties associated with the depth edge were statistically more likely. These conclusions were supported by a computational study of simulated 3D environments. Together, our results suggest that the visual system can exploit regularities in natural image and scene statistics to achieve perceptual stability at depth edges.

## Methods

All aspects of the natural scene analyses, perceptual experiments, and modeling were performed using Matlab (Mathworks, Inc.). In the perceptual experiments, stimulus presentation was controlled using the PsychToolbox-3 extensions (Brainard, 1997; Kleiner, Brainard, & Pelli, 2007; Pelli, 1997).

### Stimulus sampling

Stimuli for the scene statistics analyses and for the perceptual experiments were sampled from the natural scene dataset described in Burge et al., 2016. This dataset is composed of ∼100 stereo-images of natural scenes with precisely co-registered distance data. The stereo-images were photographed with an inter-camera separation of 6.5 cm, similar to a typical human interocular separation. The goal of the stimulus sampling was to select a set of stereo-image patches containing depth edges with relatively large monocular regions that could be (a) used to investigate the visual content and surrounding binocular context in a natural scene statistics analysis, and (b) used as stimuli in perceptual experiments.

To determine whether each pixel in a scene from the dataset was monocularly visible, we computed the ground truth horizontal disparity gradients at each image pixel directly from the distance maps (Iyer & Burge, 2018). It has been shown mathematically that if the disparity gradients—the change in disparity divided by the change in visual angle—all exceed a value of 2.0, the pixel is visible to only one eye (Bülthoff, Fahle, & Wegmann, 1991). Visual inspection of the results confirmed that this procedure was accurate on the current range maps. Stereo-image patches (3.8°x1.5°, 195×78 pixels) centered on monocularly visible points at depth edges were sampled randomly. To ensure uniqueness, we selected patches that did not overlap with each other by more than 0.2°.

We vetted the sampled patches using an automated process followed by additional manual processing. To make sure the monocular region had sufficient area for analysis, the central row in the patch was constrained to contain a set of monocularly visible points between 0.3° and 0.7° wide. In all other rows, the monocularly visible region was constrained to be at least 0.1° wide. To make sure the binocular regions were sufficiently large to analyze, the monocular region was constrained to be neighbored by at least 0.5° of relatively contiguous binocular background and foreground regions (see below for details). These criteria were chosen in order to obtain patches with monocular regions that were large enough to facilitate an informative analysis of visual patterns and binocular context (foreground/background) in natural scenes, but small enough that the neighboring binocular disparities would cause minimal diplopia during the perceptual experiments. Additionally, some patches contained local lowlights or highlights that resulted in luminance clipping. If more than 25% of the pixels either had no light recorded or were saturated, the patch was discarded. This sampling process resulted in 215 patches for analysis.

**Figure 3A & B** show two examples from the final patch set. Each example consists of a stereo-image patch and co-registered distance values at each pixel (**top and middle rows**), as well as binary masks labeling each pixel that was monocularly visible (**bottom rows**). In these binary masks, we removed minor artifacts that resulted from the range scanner as well as small regions of adjacent monocular pixels that intruded on the background. For example, if a tree or bush was visible in the background adjacent to a depth edge, there might be small speckles of monocular pixels associated with small partial occlusions. These were removed by manually editing the binary mask.

**Figure 3.**
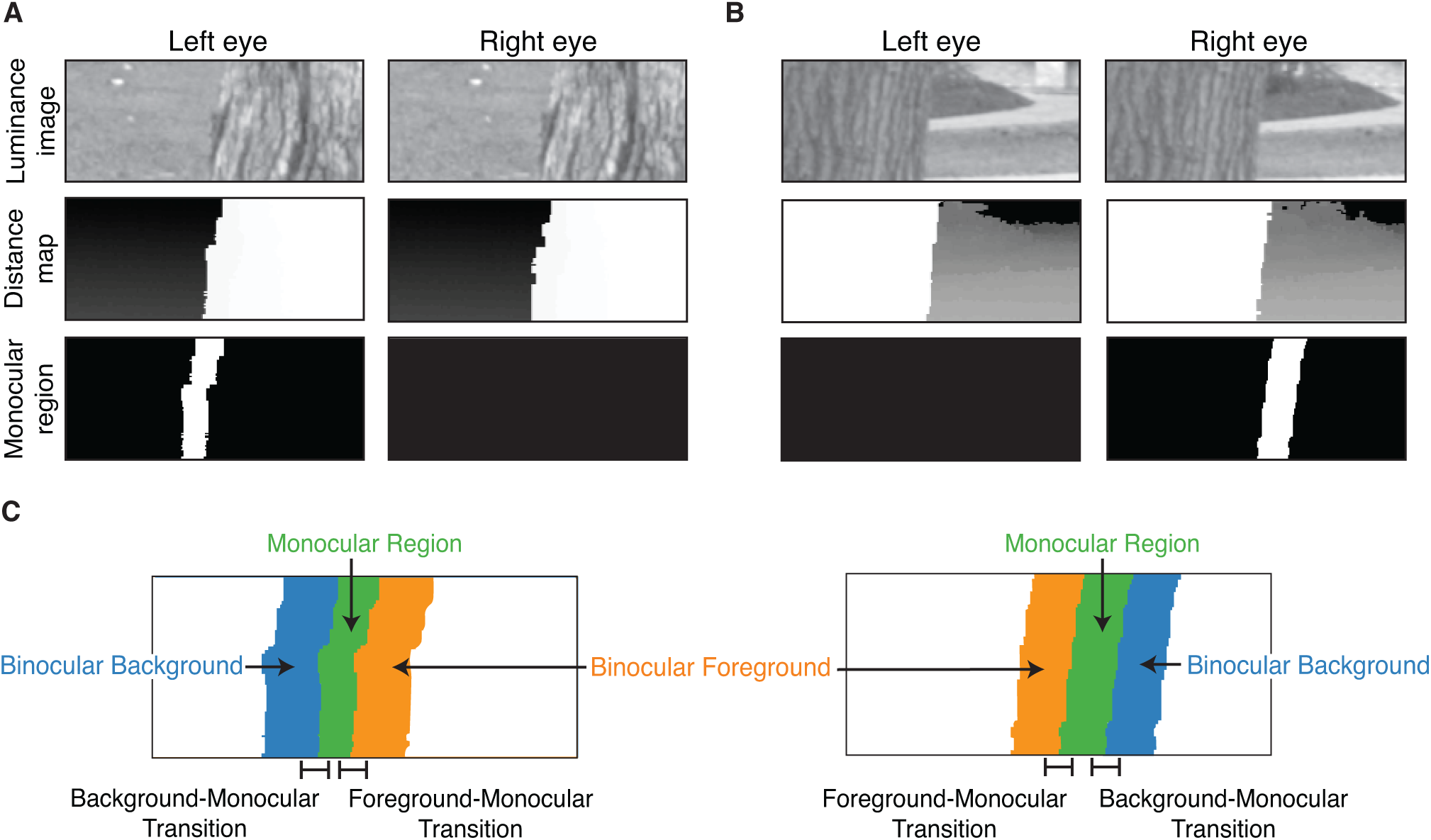
Examples of stereoscopic patches containing depth edges. **A.** Left and right eye views are shown from one of the sampled depth edges. Rows contain the luminance images (top), distance maps (middle) and monocular region binary masks (bottom). Luminance images have been histogram-adjusted for visibility. Grayscale values in the distance maps represent distance from the camera in meters, scaled to fill the full range from white (near) to black (far). Across all patches used, the distance from the cameras to the foreground region ranged from 3.1-11.1 m (average 5.4 m) and the distance from the cameras to the background ranged from 4.3-127.5 m (average 23.8). The monocular region mask indicates a single region in the left eye, consistent with the left eye seeing more of the background to the left of the tree. **B.** Same as A, but in this case the monocular region appears in the right eye. **C.** The five spatial regions of interest used for analysis are illustrated for both patches.

### Natural Scene Statistics

To organize our analysis of the statistics of depth edges in natural scenes, we defined five regions of interest (**Figure 3C**). These regions were the monocular region (green), the adjacent binocular foreground (orange), the adjacent binocular background (blue), and two transition regions from the binocular background to the monocular region and from the binocular foreground to the monocular region (black lines). The binocular regions consisted of the 0.5° wide image areas neighboring the monocular region to the left or right. The transition regions contained 0.4° wide image areas centered on the transition between the monocular region and the binocular background or foreground regions. The forthcoming results are similar for other similar region sizes.

From the distance maps, we calculated the mean distance of the surfaces in the monocular region, the adjacent binocular foreground region, and the adjacent binocular background region. From the images, we computed the mean luminance and the mean contrast (i.e. square root of the mean squared luminance deviation) within each of these regions as well. In the binocular-monocular transition regions, we focused on analyzing the changes in visual appearance associated with transitions between surfaces. For example, a large horizontal luminance derivative would reflect a strong vertical edge, suggesting a transition between two surfaces with different patterns. To quantify the strength of the vertical luminance edge between the monocular region and adjacent binocular regions, we used 5-tap derivative filters to compute the mean magnitude of the horizontal luminance derivative (Farid & Simoncelli, 1997). As a control measure, we repeated this analysis for vertical derivatives, which are not expected to be particularly strong at horizontal surface transitions.

Prior to statistical analyses, all of the image-based measurements described above were normalized. Specifically, the mean luminance within each region and the mean magnitude of luminance derivatives within each transition region were normalized by the mean luminance of the entire patch. The mean contrast within each region was normalized by the mean contrast of the entire patch. Two-tailed Wilcoxon signed rank tests were used to examine statistically significant differences in the visual properties of different regions. Non-parametric effect sizes for these tests were calculated as 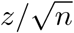, where *z* refers to the *z* score of the Wilcoxon test statistic and *n* refers to the number of differences used to calculate the test statistic (Fritz, Morris, & Richler, 2012).

It is important to note that the natural scene dataset used here does not contain objects that are closer than 3 m. This limitation of the dataset should be considered when interpreting the results of our analysis because prior work suggests that the prevalence of monocularly visible regions depends on the distribution of object distances in a scene (Langer & Mannan, 2012). That said, we later report the results of a geometric simulation that parametrically examined how object distance affects the geometric causes of monocular regions (see Results). The findings from the simulation suggest that the main conclusions of this natural scene analysis are likely to hold not just for surfaces that are far away, but also for surfaces that are up close.

### Perceptual Experiments

Two perceptual experiments were conducted to examine how the visual properties of natural depth edges affect perceptual stability. Experiment 1 was conducted in an exploratory manner and used to develop a set of hypotheses and appropriate statistical tests. Experiment 2 tested these hypotheses with a second set of observers, a larger, more diverse set of stimuli and a higher dynamic range display. We first describe the methods in the two experiments that were the same, and then indicate the differences between them.

### General Methods

#### Observers

All observers had normal or corrected-to-normal visual acuity and normal stereoacuity, as determined by the Randot stereo test (detection of 70 arcsecs or less). Observers gave informed consent prior to starting the experiment. The procedures were approved by the institutional review board at the University of California, Berkeley and were consistent with the Declaration of Helsinki. Experiment 1 had 10 observers (5 female, mean age = 31.0□2.6 years) and Experiment 2 had 15 observers (9 female, mean age = 27.7□5.4 years).

#### Stimulus Presentation

Patches from the dataset described above were presented to observers stereoscopically on gamma-linearized displays. Observers were positioned via a forehead and chin rest at a viewing distance of approximately 60cm (1 pixel subtended ∼0.02-0.03°). The stimulus was designed to simulate viewing a portion of a natural scene through a window (**Figure 4**). Specifically, stereoscopic scene patches were viewed through a virtual window that was specified by disparity to be at the display distance. The window subtended the same visual area as each patch (3.75°x1.5°). Using recently developed techniques described in Iyer & Burge (2018), left and right eye image patches were cropped at corresponding points such that the depth edge had five arcminutes of uncrossed disparity with respect to the screen. This procedure ensured that all depicted surfaces in the scene were stereoscopically defined to be behind the plane of the display and the frame of the window.

**Figure 4.**
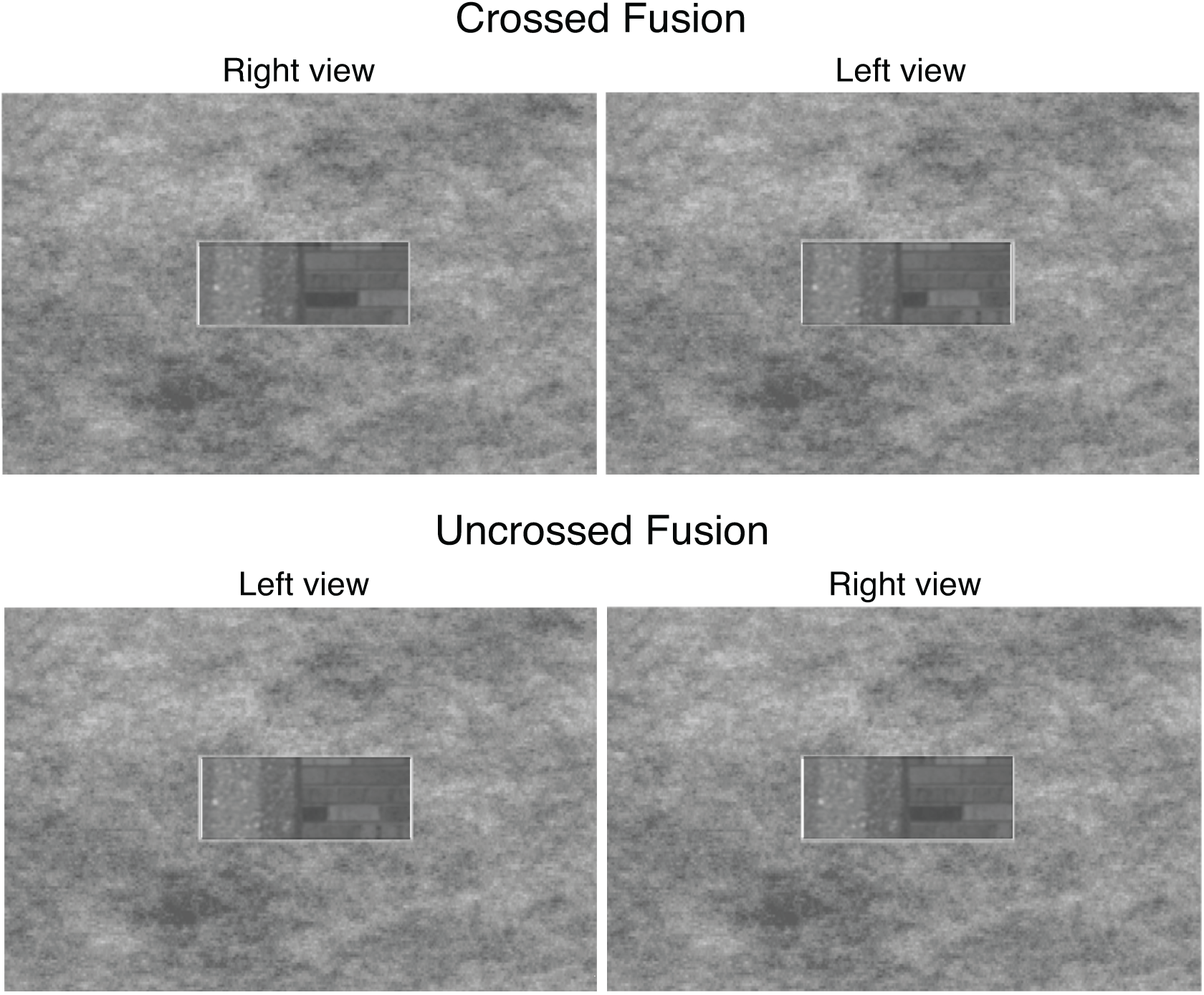
Example stimulus. Left and right eye views are shown (oriented for crossed fusion in the upper row, and uncrossed fusion in the lower row). Each stereoscopic patch was placed in a window surrounded by 1/f noise, on an uncrossed disparity pedestal so that all content appeared behind the window frame. Stimulus is shown as presented in Experiment 1, and the surrounding background is cropped for visibility. Images have been histogram adjusted for visibility.

On each trial, observers viewed a patch for 5 seconds through the virtual window, and were then asked on a separate response screen to evaluate the visual stability of the patch using a continuous slider response. To explain the task, they were shown an example of a rivalrous stimulus and given descriptions of types of visual instability (e.g., shimmering or alternating between two states). Each response was recorded on a 100-point scale, with 100 indicating the most unstable. We refer to these responses as *instability ratings*. Pilot testing confirmed that even if there was local instability at the depth edge, observers could fuse these stimuli as a whole and experience depth percepts of the foreground and background (White & Burge, 2019). For the current study we chose a visual stability rating task, because previous work suggests that stability is a salient aspect of perception when binocular fusion fails (Hoffman & Banks, 2010; Shimojo & Nakayama, 1990). All observers first completed an initial practice session in which they observed and rated example patches. For Experiment 2, this practice session included all stimuli in order to facilitate better response reliability during the main task. Data from the practice session were not included in analysis. Following the practice session, observers completed two repetitions per patch. Patches were not repeated until all stimuli had been seen once. We calculated the test-retest reliability via the squared Pearson correlation coefficients (r^2^) between each observer’s ratings from the 1^st^ and 2^nd^ presentations of each patch. Observers with r^2^ values below 0.1 were excluded from further analysis (3 observers in Experiment 1, and none in Experiment 2).

### Experiment 1

#### Stimuli

We selected a subset (n = 92) of the patches from the natural scene statistics analysis for an exploratory study. To reduce the number of features that varied across different patches, the mean luminance and contrast of each patch were fixed prior to displaying them to the observers. The mean luminance of each patch was set to 30% of the maximum luminance output of the display. The normalized root-mean-squared (RMS) contrast of each patch when presented onscreen was fixed to 0.25. Note that the luminance and contrast within and across the regions of interest still varied in the patches – fixing the global mean luminance and global mean contrast simply allowed us to concentrate our analyses on the impact of local image properties. Stimuli were presented stereoscopically on a liquid crystal display (LCD) (ViewSonic V3D231, 50.9 x 28.6cm, 1920 x 1080 pixels), which supports passive stereoscopic presentation via polarizing filters. Observers wore polarizing glasses so that stimuli could be viewed dichoptically (specifically, alternating rows of pixels on the screen were visible to each eye). In this experiment, the viewing window was surrounded by a 1/f noise field. The maximum luminance through the filters was approximately 63 cd/m^2^, and stimulus settings were selected to eliminate any visible crosstalk.

### Experiment 2

#### Stimuli

In this second experiment, we used a larger set of patches to obtain a more diverse set of stimuli (n = 125). In addition, we did not fix the mean luminance or mean contrast of the stimuli, as we did in Experiment 1. By allowing these values to vary, we could test whether the findings from Experiment 1 generalized to more diverse stimuli (i.e., they are not specific to the luminance and contrast values selected for that experiment). The results were highly similar across experiments. Stimuli were presented stereoscopically on a mirror haploscope with two LCDs to provide a higher luminance range (LG 32UD99-W, 69.6 x 39.3 cm, 1920 x 1080 pixels, maximum luminance of approximately 330 cd/m^2^). The stimulus was adjusted so that the vergence demand of the virtual window was equal to the monitor distance. The virtual window was surrounded by a 1/f noise field, as in Experiment 1, but the texture was removed from the area immediately around the window in order to reduce visual clutter.

### Modeling instability ratings

We used the same statistical analysis in both experiments. First, we defined a set of image features, the values of which varied across the stimuli. Then, we defined a set of linear mixed models that used these features as predictors for patch instability ratings. In each model, different observers were modeled with random intercepts, and the usefulness of different features for predicting the responses was examined (see Results for details). Initial analyses of Experiment 1 were conducted in an exploratory manner based on the results of the scene statistics analysis. Based on these exploratory analyses, we hypothesized that the strength of the vertical edges at the binocular-monocular transition regions modulated the perceptual stability of the depth edges. We then conducted a second experiment to test this hypothesis on a larger independent sample of perceptual data.

In a follow up analysis, we asked whether higher contrast in the patches was associated with higher perceptual instability, based on predictions from previous work (L. Liu, Tyler, & Schor, 1992). For each observer, we converted instability ratings to z-scores by subtracting the mean response and dividing by the response standard deviation. We calculated Pearson correlation coefficients between the contrast in each region and the mean z-score for each patch. For correlation values, 95% confidence intervals are calculated via bootstrapping.

## Results

### Natural Scene Statistics

We analyzed the statistical regularities of scene and image properties in the immediate vicinity of depth edges in natural scenes, with an emphasis on the transitions between monocular and binocular regions. If robust regularities are present, the visual system might exploit these regularities to determine which regions of the scene are monocularly and binocularly visible.

First, we analyzed the difference between the mean distance to the surfaces in the monocularly visible region of the scene and the mean distances to the surfaces in the adjacent binocularly visible background and foreground regions. This analysis was possible because each image in the dataset had laser-based distance measurements co-registered to each image pixel (Burge et al, 2016). When a monocular region primarily consists of the background occluded by the foreground (**Figure 2A**), the distance of monocularly visible points from the camera, on average, should be more similar to the distance of the binocular background than the distance of the binocular foreground. Conversely, when a monocular region primarily consists of a self-occlusion (**Figure 2B**), the distance of the monocularly visible points, on average, should be more similar to the binocular foreground. We found that the distance of surfaces in the monocular regions tended to be substantially more similar to the adjacent binocular background than the foreground. **Figure 5A** shows histograms of the distance between each monocular region and the adjacent binocular foreground (orange) and the adjacent binocular background (blue) – that is, the mean distance in the monocular region minus the mean distance in the adjacent binocular regions. A value of zero indicates that points in the monocular region had the same average distance as points in the relevant binocular region. Positive and negative values indicate that the distance of points in the monocular region was farther and closer, respectively, than the distance of points in the binocular region. The median absolute distance of points in the monocular region from the binocular foreground was 9.59 m, while the median absolute distance of points in the monocular region from the binocular background was 0.10 m. The absolute distance of points in the monocular region to points in the binocular background was significantly smaller than the distances to points in the binocular foreground (*p* << 0.001, *z* = 12.71, *effect size* = 0.87). This result suggests that, in this dataset, the vast majority of monocular regions were caused primarily by background occlusions.

**Figure 5.**
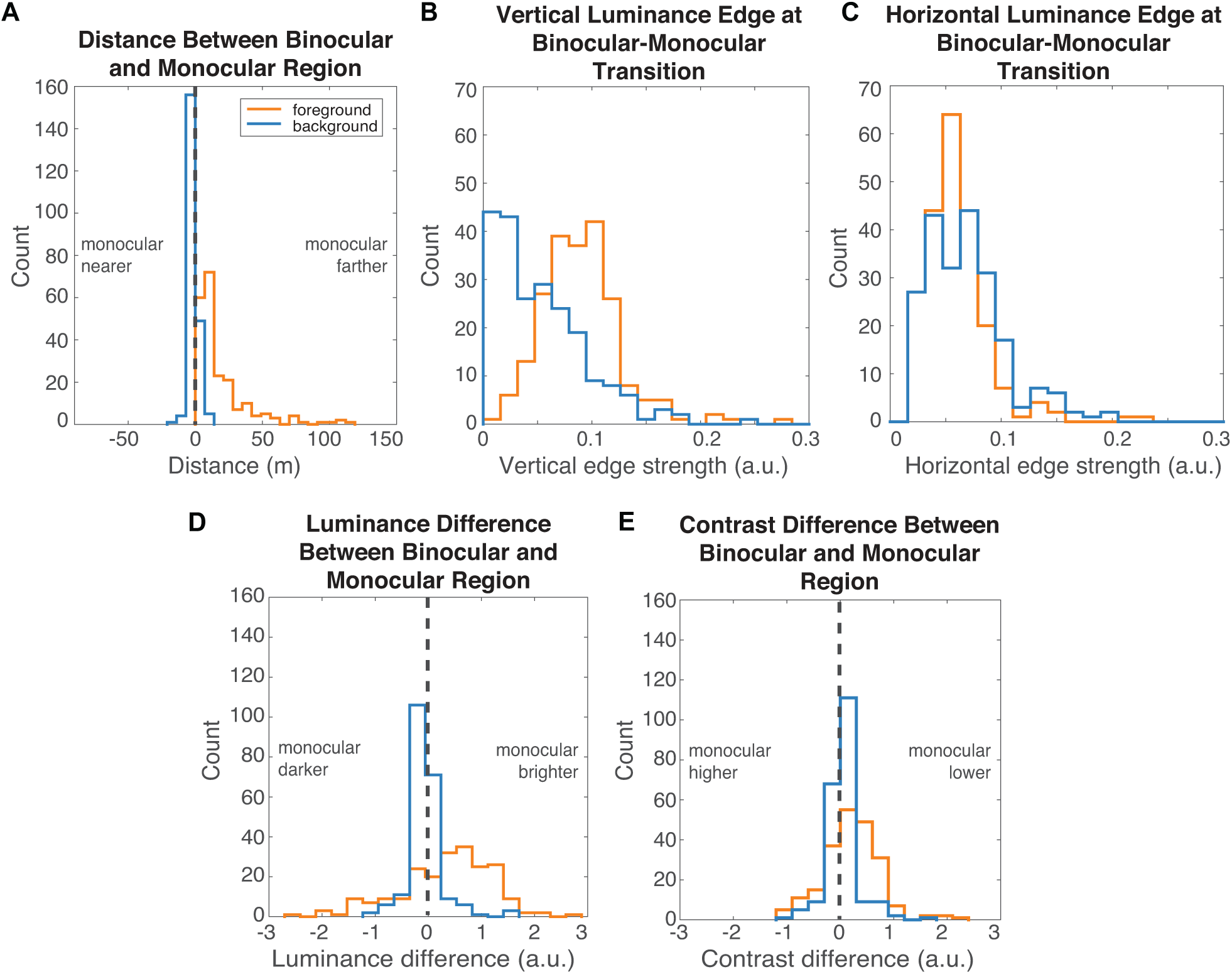
Properties of monocular regions and the surrounding binocular context. Histograms show the frequency of: (A) the distances between binocular foregrounds/backgrounds and monocular regions within a patch, (B) the vertical luminance edge strength at the transitions from binocular foregrounds/backgrounds to monocular regions, (C) the horizontal luminance edge strength at the transitions from binocular foregrounds/backgrounds to monocular regions, (D) the differences in luminance between binocular foregrounds/backgrounds and monocular regions, and (E) the differences in contrast between binocular foregrounds/backgrounds and monocular regions. Distance was calculated in units of meters. Vertical and horizontal edge strengths represent the normalized magnitude of the derivative filters in the respective transition regions (i.e., foreground-monocular and background-monocular). Luminance is represented in arbitrary units, and contrast is represented as normalized RMS pixel luminance. A.u. indicates arbitrary units.

We reasoned that if a monocular region associated with a depth edge is caused by a background occlusion (i.e., is an extension of the neighboring binocular background), then there should be reliable information about this fact in the luminance pattern within the *transition regions* (see **Figure 3C**) (Y. Liu, Cormack, & Bovik, 2009). Specifically, when a monocular region is an extension of the neighboring binocular background there should be a relatively strong vertical luminance edge at the foreground-monocular transition region, because this will coincide with the depth edge. Relatedly, there should be a relatively weak vertical edge at the background-monocular transition region, because the binocular background and the monocular region are likely to contain the same surfaces. On the other hand, if the foreground surface occludes itself— a self-occlusion **(Figure 2B)**—there will be no depth edge in the foreground-monocular transition region, and there should therefore be a relatively weak vertical luminance edge in the foreground-monocular transition region. A stronger luminance edge is thus expected in the background-monocular transition region (**Figure 2B**). Note that these inferences are based on the assumption that background and foreground surfaces have visually distinct textures, and that the statistics of these textures are relatively stationary within a surface. These assumptions have not been previously tested empirically.

The current data suggest that these assumptions are sufficiently accurate for natural image patches to meet these expectations. The vertical luminance edge strength (calculated as described in the Methods) in the foreground-monocular transition region tended to be higher (0.09) than at the vertical luminance edge strength in the background-monocular transition (0.05) (**Figure 5B**; *p* << 0.001, *z* = 10.7, *effect size* = 0.73). If this pattern is tied to the surface in the monocular region, we hypothesized that this pattern should be *specific* to vertical luminance edges, because edges that are vertical indicate a horizontal transition between surfaces. To test this hypothesis, we repeated this analysis for horizontal luminance edge strength. With horizontal luminance edges, we found a small but statistically significant difference in the opposite direction. The horizontal edge strength at the foreground-monocular transition was slightly but significantly lower (0.057) than the horizontal edge strength at the background-monocular transition (0.064) (**Figure 5C**; *p* << 0.001, *z* = 3.43, *effect size* = 0.23). We speculate that this effect occurs because the abrupt transition between surfaces at the foreground-monocular transition precludes any strong horizontal continuity. Importantly, with these analyses, we cannot rule out the possibility that some of these monocular regions were caused by hybrid occlusions: a background occlusion combined with a thin self-occlusion at the curved edge of a foreground object, for example. Such hybrid occlusions would still provide a vertical luminance edge within the defined transition region. In the forthcoming geometric simulation, we examine the prevalence of hybrid occlusions directly.

To further test the conclusions of the transition analysis (i.e., that monocular regions tend to primarily belong to the binocular background, and therefore share similar visual features), we examined the luminance and contrast within the monocular regions relative to the binocular regions (**Figure 5D and E**). Consistent with the previous analysis, the results showed that these visual properties were more similar to the binocular background than to the binocular foreground. Specifically, the median absolute luminance difference from the background (0.08) was approximately ten times lower than the median difference from the foreground (0.74) (*p* << 0.001, *z* = 11.6, *effect size* = 0.79). Recall that luminance is represented in arbitrary linear units. The median absolute contrast difference between the monocular region and the background (0.10) was more than three times lower than the median absolute contrast difference between the monocular region and the foreground (0.36) (*p* << 0.001, *z* = 8.95, *effect size* = 0.61). Indeed, the strength of the vertical luminance edge between two regions was highly correlated with both the luminance difference (*r* = 0.57, 95% confidence interval 0.49-0.64, *p* << 0.001) and the contrast difference (*r* = 0.51, 95% confidence interval 0.42-0.61, *p* << 0.001). We speculate that these measures are highly correlated because they all reflect the same underlying image structure in the neighborhood of depth edges.

Collectively, these natural scene analyses suggest that monocularly visible regions in natural scenes are associated with robust statistical regularities. The results suggest that monocular regions in natural scenes are more likely to be composed primarily of an occluded extension of the binocularly visible background than an occluded portion of the binocularly visible foreground, at least within scenes that are similar to those in this particular dataset (e.g., relatively far object distances, relatively sparse object density). According, the monocular regions we analyzed tended to visually similar to the adjacent binocular background. In particular, the transition from the foreground to the monocular region was associated with a strong vertical luminance edge, whereas the transition from the monocular region to the background was not. If the visual system is adapted to these statistical regularities in the natural environment, the visual similarity of the monocular region with the background, and perhaps the visual dissimilarity of the monocular region with the foreground, should be associated with increased perceptual stability of depth edges in natural scenes. We similarly predict that greater perceptual instability will result from stimuli in which these statistical regularities in the luminance images are violated. To test these predictions, we performed a pair of perceptual experiments.

### Perceptual Experiments

As a reminder, we conducted two perceptual experiments in which observers viewed a subset of the natural stereo-image patches from the previous analysis and rated the perceived instability of the depth edge. The perceptual experiments were designed to determine whether the statistical regularities associated with depth edges in natural scenes predict perceptual instability. Before examining this question, we first established the consistency of ratings within and across observers (recall that each observer rated each patch twice). The mean correlation of the observer ratings from the first and second stimulus presentations was 0.45 in Experiment 1 and 0.61 in Experiment 2, suggesting reasonable consistency within observers. To obtain mean instability ratings for each patch, we thus averaged the first and second ratings of each observer. **Figure 6** shows the mean and standard deviation of these z-scored ratings across observers for each patch, ranked from highest to lowest. Across patches, the perceived stability varied considerably. The standard deviation of instability ratings across observers was relatively consistent across patches and was also relatively small compared to the change in stability across the patch ensemble. Together, these results suggest that meaningful comparisons of perceived stability can be made across this stimulus ensemble.

**Figure 6.**
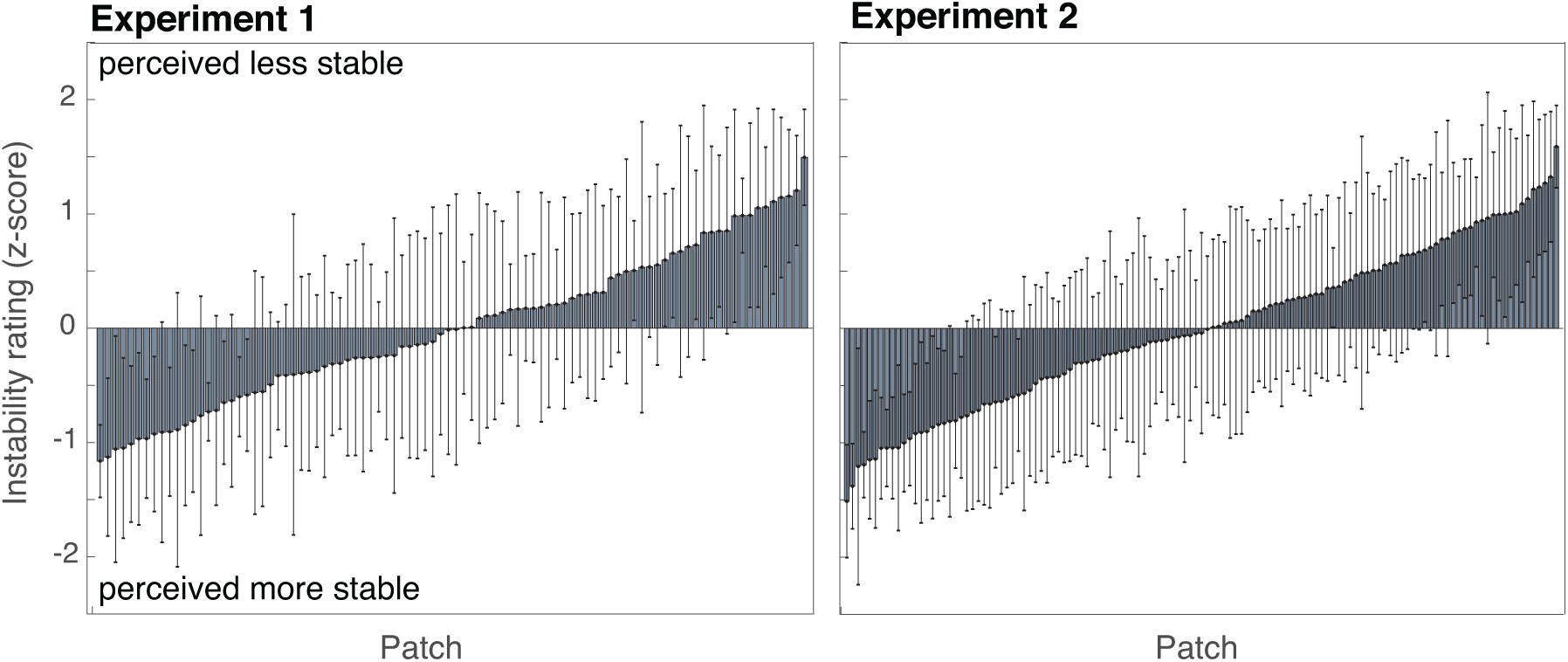
Instability ratings for each natural stereoscopic patch stimulus in both Experiments. Each bar represents the mean z-score for a patch across observers, ordered from lowest to highest. Error bars represent the standard deviation of the mean.

Now, we turn to an analysis of the factors that may contribute to perceptual instability. Before examining the impact of image-based properties on perceptual instability, we first determined how much of the variation in perceived instability could be attributed to the relative disparity between the foreground and background surfaces. Depth edges with a large binocular disparity between the foreground and background may tend to appear unstable when they exceed Panum’s fusional area, for the simple reason that they are difficult to fuse. With large disparities, the background is likely to appear diplopic when the foreground is fused and vice versa. In the stimulus set, the magnitude of the depth steps across the edges corresponded to relative disparities ranging from 0.3-0.8°. We found that relative disparity was significantly correlated with the mean instability ratings in both experiments (Experiment 1: *r* = 0.54, 95% confidence interval 0.32-0.70, *p* << 0.001; Experiment 2: *r* = 0.59, 95% confidence interval 0.47-0.70, *p* << 0.001), accounting for 29% and 34% of the total variance in mean instability ratings, respectively. Hence, as expected, larger relative disparities are associated with more perceptual instability.

To model the effects of the image-based properties on instability ratings, we fit the raw response data with a set of mixed linear regression models. Based on the analyses of natural scene statistics, we formulated two nested models: the *Disparity-Only model* and the *Edge model*. The Disparity-Only model included the magnitude of relative disparity as the only predictor. The Edge model included the vertical luminance edge strength in each of the two binocular-monocular transition regions as two additional predictors. We chose these two predictors because the natural scene statistics analysis suggested that a strong vertical luminance edge at the foreground-monocular transition is likely in natural depth edges. We thus hypothesized that strong foreground-monocular vertical edges should be associated with higher perceptual stability, but that background-monocular vertical edges should not.

The results suggest that higher perceptual stability is associated with image patterns that are more likely given the natural image and scene statistics. **Table 1** summarizes these results for each experiment. The observed versus the fitted ratings are plotted in **Figure 7**. The Disparity-Only model and the Edge model both indicate a significant effect of disparity magnitude. In the Edge Model, vertical luminance edge strength in the binocular-monocular transition region was also a significant predictor of the ratings: stronger vertical edges in the foreground-monocular transition region were associated with greater perceptual stability (lower ratings) and stronger vertical edges in the background-monocular transition region were associated with greater perceptual *instability* (higher ratings). A vertical edge at an atypical location relative to the monocular region may cause the visual system to misinterpret monocular regions as being binocular, and the increased instability may thus result because the visual system tries and fails to fuse monocular regions with binocular regions.

**Table 1.**
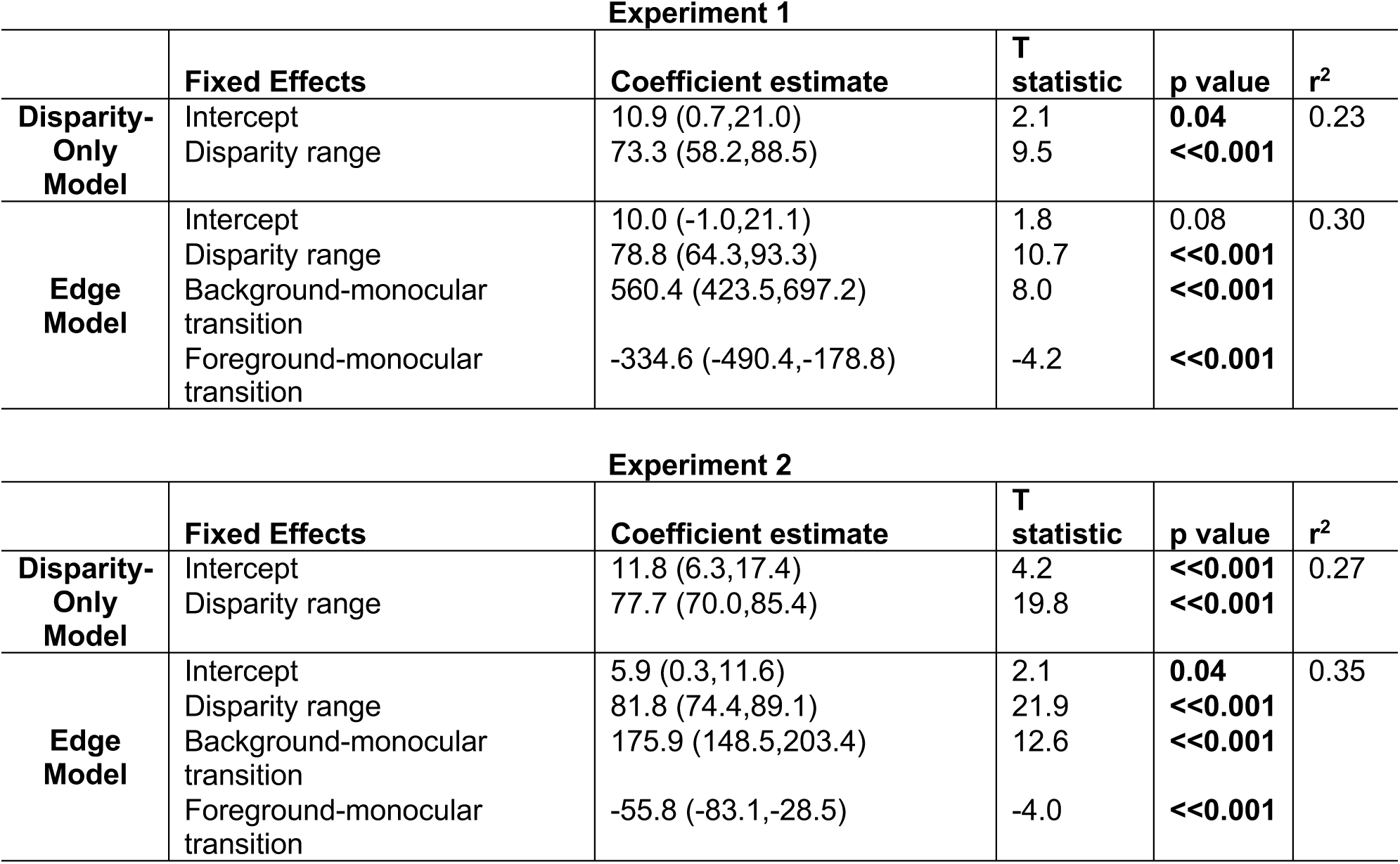
Regression model results. For each experiment, results are shown for the two nested models: the Disparity-Only model and the Edge model. Confidence intervals (95%) on each of the coefficient estimates are indicated in parentheses. T-statistics were used to assess whether these estimates were significantly different from zero. Significance values less than 0.05 are bolded. Random intercepts were fit per observer in each experiment, with a standard deviation of 9.1 and 8.0 in Experiments 1 and 2, respectively.

| Experiment 1 |  |  |  |  |  |
| --- | --- | --- | --- | --- | --- |
|  | Fixed Effects | Coefficient estimate | T statistic | p value | r<sup>2</sup> |
| Disparity-Only Model | Intercept | 10.9 (0.7,21.0) | 2.1 | 0.04 | 0.23 |
|  | Disparity range | 73.3 (58.2,88.5) | 9.5 | <<0.001 |  |
| Edge Model | Intercept | 10.0 (-1.0,21.1) | 1.8 | 0.08 | 0.30 |
|  | Disparity range | 78.8 (64.3,93.3) | 10.7 | <<0.001 |  |
|  | Background-monocular transition | 560.4 (423.5,697.2) | 8.0 | <<0.001 |  |
|  | Foreground-monocular transition | -334.6 (-490.4,-178.8) | -4.2 | <<0.001 |  |

| Experiment 2 |  |  |  |  |  |
| --- | --- | --- | --- | --- | --- |
|  | Fixed Effects | Coefficient estimate | T statistic | p value | r<sup>2</sup> |
| Disparity-Only Model | Intercept | 11.8 (6.3,17.4) | 4.2 | <<0.001 | 0.27 |
|  | Disparity range | 77.7 (70.0,85.4) | 19.8 | <<0.001 |  |
| Edge Model | Intercept | 5.9 (0.3,11.6) | 2.1 | 0.04 | 0.35 |
|  | Disparity range | 81.8 (74.4,89.1) | 21.9 | <<0.001 |  |
|  | Background-monocular transition | 175.9 (148.5,203.4) | 12.6 | <<0.001 |  |
|  | Foreground-monocular transition | -55.8 (-83.1,-28.5) | -4.0 | <<0.001 |  |

**Figure 7.**
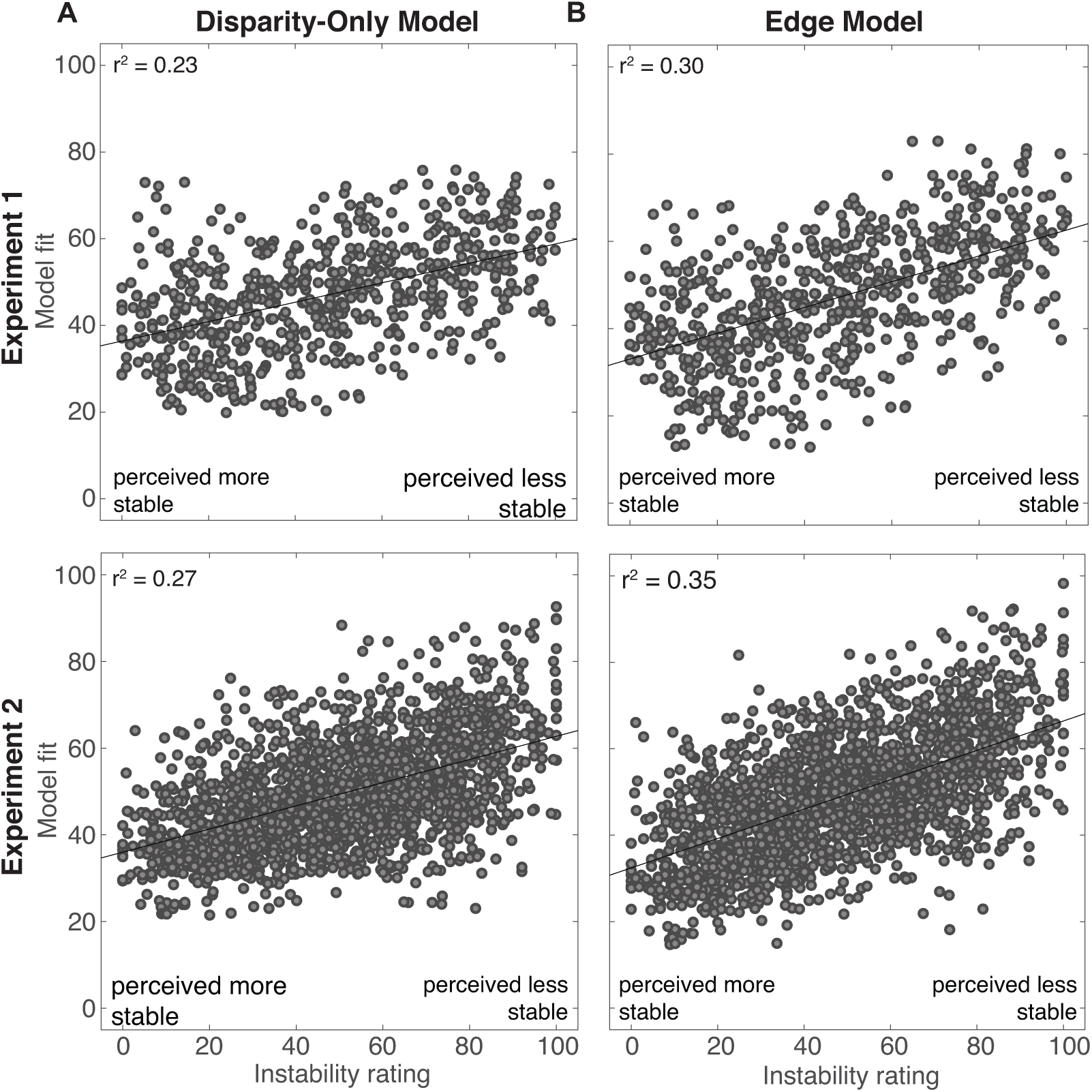
Actual versus fitted instability ratings for the Disparity-Only model (A) and Edge model (B). Each panel shows the instability ratings of all observers on the abscissa, and the ratings produced by the best fit model on the ordinate. The top row shows the data from Experiment 1, and the bottom row shows the data from Experiment 2.

A likelihood-ratio test indicated that the Edge Model was a significantly better fit to the data (Experiment 1: *LR* = 61.61, □ *df* = 2, *p* << 0.001; Experiment 2: *LR* = 194.52, □ *df* = 2, *p* << 0.001). This result was supported by a comparison of each model’s Akaike Information Criterion (AIC), which accounts for the increased number of parameters in the Edge Model (Experiment 1: Disparity-Only Model = 5922.9, Edge Model = 5892.1; Experiment 2: Disparity-Only Model = 16649, Edge Model = 16458). We also computed the proportion of the response variance explained by the Edge Model as compared to the Disparity-Only Model. The Edge Model explained 30% and 35% of the response variance in Experiments 1 and 2, while the Disparity-Only model explained 23% and 27%. These percentages should be interpreted relative to the expected amount of explainable variance in the modeled data, which is limited by the variability in repeated presentations of the same stimulus to the same observer. Using the method described in Hsu, Borst, & Theunissen (2004), we estimated that the explainable variance in the mean responses in the first experiment was 50% and in the second experiment was 62% (see also Holdgraf et al., 2017). These results are approximate, because we only had two repeats per observer, but we can conclude that the Edge Model captures a reasonable portion of the explainable variance within and across observers in the current data. Thus, the results of this analysis suggest that image cues at the binocular-monocular transition regions of a depth edge affect perceptual stability, and that this effect is consistent with observed statistical regularities in real depth edges from natural scenes.

To further explore this result, we examined to what extent luminance or contrast differences between the monocular and binocular regions contributed to the mediating effects of transition region vertical edges on perceptual stability. We created two additional models, each with two predictors in addition to disparity magnitude: the *Luminance model* and the *Contrast model*. In the Luminance model, we used the luminance difference between the monocular region and the background, and between the monocular region and the foreground (**Figure 5D**) as predictors. In the Contrast model, we replaced these with the contrast differences (**Figure 5E**). Both of these signals can contribute to the strength of the vertical edge. The results suggest that the luminance differences, and to some extent contrast differences, contributed to the influence of binocular-monocular vertical edges on perceptual stability. Specifically, greater luminance differences between the monocular region and the binocular foreground were associated with more stability (lower ratings), and greater luminance differences between the monocular region and the binocular background were associated with more instability (higher ratings). Greater contrast differences between the monocular region and the binocular background were associated with more instability (this effect was only statistically significant in Experiment 2). The full results are reported in **Table 2**. In practice, the identification of binocular corresponding and non-corresponding points is likely to also rely on higher order image features. However, these results provide some insight into the nature of the surface differences that may facilitate binocular fusion.

**Table 2.** Regression model results for luminance and contrast differences. Data are formatted the same as in Table 1.

| Experiment 1 |  |  |  |  |  |
| --- | --- | --- | --- | --- | --- |
|  | Fixed Effects | Coefficient estimate | T statistic | p value | r<sup>2</sup> |
| Luminance Model | Intercept | 23.53 (12.60,34.45) | 4.23 | <<0.001 | 0.31 |
|  | Disparity range | 71.78 (57.34,86.22) | 9.76 | <<0.001 |  |
|  | Background-monocular difference | 37.10 (17.93,56.28) | 3.80 | <<0.001 |  |
|  | Foreground-monocular difference | -42.63 (-53.78,-31.49) | -7.51 | <<0.001 |  |
| Contrast Model | Intercept | 7.46 (-3.20,18.12) | 1.37 | 0.17 | 0.24 |
|  | Disparity range | 76.71 (61.36,92.06) | 9.81 | <<0.001 |  |
|  | Background-monocular difference | 8.00 (-0.43,16.43) | 1.86 | 0.06 |  |
|  | Foreground-monocular difference | 1.45 (-4.16,7.05) | 0.51 | 0.61 |  |

| Experiment 2 |  |  |  |  |  |
| --- | --- | --- | --- | --- | --- |
|  | Fixed Effects | Coefficient estimate | T statistic | p value | r<sup>2</sup> |
| Luminance Model | Intercept | 15.51 (9.80,21.22) | 5.33 | <<0.001 | 0.29 |
|  | Disparity range | 74.97 (67.37,82.57) | 19.35 | <<0.001 |  |
|  | Background-monocular difference | 9.98 (6.64,13.32) | 5.86 | <<0.001 |  |
|  | Foreground-monocular difference | -5.50 (-7.22, -3.78) | -6.27 | <<0.001 |  |
| Contrast Model | Intercept | 10.21 (4.55,15.88) | 3.53 | <<0.001 | 0.28 |
|  | Disparity range | 78.27 (70.60,85.94) | 20.02 | <<0.001 |  |
|  | Background-monocular difference | 5.74 (1.96,9.52) | 2.98 | 0.003 |  |
|  | Foreground-monocular difference | 0.64 (1.97,3.25) | 0.48 | 0.63 |  |

In a final analysis, we examined how perceptual instability experienced by our observers when viewing natural depth edges relates to stimulus features that are known to exacerbate binocular rivalry in simple stimuli. Previous studies on binocular rivalry that used synthetically mismatched stimuli (e.g., sinusoidal gratings with different orientations in the two eyes), have reported that low contrast stimuli tend to be more perceptually stable than their high contrast counterparts. Sometimes, stimuli that cause unstable percepts at high contrast are fused into stable unified percepts at low contrast. For example, when orthogonal gratings are dichoptically presented at high contrast, they cause strikingly unstable percepts that alternate between the two grating orientations. When the same orthogonal gratings are presented at low contrast, they are perceived as a stable plaid (L. Liu et al., 1992). Does the relationship between contrast and perceptual instability hold in natural images when monocular regions are caused by depth edges? First, we examined the correlation between the instability ratings and the image contrast in the binocular background, monocular, and binocular foreground regions in Experiment 2. As expected, binocular instability tended to be higher when contrast was higher. Higher instability ratings were associated with higher contrast in the monocular region (*r* = 0.20, 95% confidence interval 0.07-0.33, *p* = 0.02) and with higher contrast in the adjacent binocular foreground (*r* = 0.31, 95% confidence interval 0.15-0.45, *p* << 0.001). This relationship was weaker and not statistically significant in the binocular background (*r* = 0.17, 95% confidence interval 0.04-0.30, *p* = 0.06). Next, we examined the correlation between instability ratings and contrast in the transition regions. Instability ratings were higher with higher contrast in the background-monocular transition region (*r* = 0.19, 95% confidence interval 0.05-0.32, *p* = 0.03), but the same was not true for the foreground-monocular transition region (*r* = −0.097, 95% confidence interval −0.25-0.09, *p* = 0.28). The absence of a correlation between perceptual instability and contrast in the foreground-monocular transition region may be connected to the natural scene statistics. Because *dissimilarity* between the monocular region and the binocular foreground is highly likely, a high contrast in this transition region provides information about the existence of an edge in the natural environment, which may facilitate rather than impede fusion.

Taken together, these results suggest that depth edges with features that are statistically more likely are also likely to be more perceptually stable. Because monocular regions tend to be similar to the adjacent binocular background than to the adjacent binocular foreground in natural scenes, the visual system might be inclined to correctly identify these regions as monocular when they share features that are similar to the binocular background instead of the foreground.

### Geometric Simulation

The results from the natural scene statistics analysis suggest that natural depth edges have robust statistical regularities that could be exploited by the visual system (i.e., monocularly visible regions tend to share visual properties with the binocular background). The perceptual studies suggest that stimuli adhering to this statistical regularity are more likely to be perceived as stable, and stimuli that violate it are more likely to be perceived as unstable. One potential caveat with these results is that the sampled patches used in this study were drawn from a set of outdoor scenes that were photographed such that the closest objects were 3m away. If the results depend on viewing distance, these may not generalize to near viewing distances. In scenes with close by objects, it is possible that self-occlusions and hidden surfaces may occur more frequently than in the scenes that we analyzed. If so, the conclusions that we have drawn may apply only to distant visual scenes.

Prior work has used geometric modeling as an additional tool to explore potential statistical regularities in binocular and monocular visibility in 3D scenes (Hansard, 2012; Langer & Mannan, 2012; Langer et al., 2016). This approach is appealing because it allows the modeler to parametrically vary properties of a scene (e.g., object distance and clutter). For example, Langer & colleagues asked how the probability of points that are visible only monocularly varies as a function of object distance (Langer & Mannan, 2012; Langer et al., 2016). They found that the probability that a given point is monocularly visible increases monotonically as a function of its distance from the viewer, and that this increase in probability is more rapid when the objects are smaller. However, because the simulated scenes were populated only with identical planar surfaces, self-occlusions were not present in these simulations. Thus, the relative frequency of the different causes of monocular regions (the topic of the current investigation) could not be examined. Building on this prior work, we performed a parametric study of simulated 3D scenes populated with objects capable of self-occlusion. Specifically, we examined how changes in viewing distance and the properties of the 3D environment change the proportion of monocular regions tend to be due to background occlusions, self-occlusions, or hidden surfaces. This simulation also allowed us to explore the prevalence of hybrid occlusions that contain multiple monocularly visible surfaces.

Each simulation included a binocular observer positioned within a different environment. Each environment was parameterized in terms of its scale (i.e. maximum object distance), its density (i.e. total number of objects), and the size of the objects within it (as a proportion of the scale). Objects were defined as polygons with a random number of sides (up to 7), and were positioned randomly within the virtual observer’s central visual field. Ray tracing was used to determine whether each visible scene point was binocularly or monocularly visible. We limited our analyses to the horizontal plane of regard. **Figure 8A** illustrates two example simulations.

**Figure 8.**
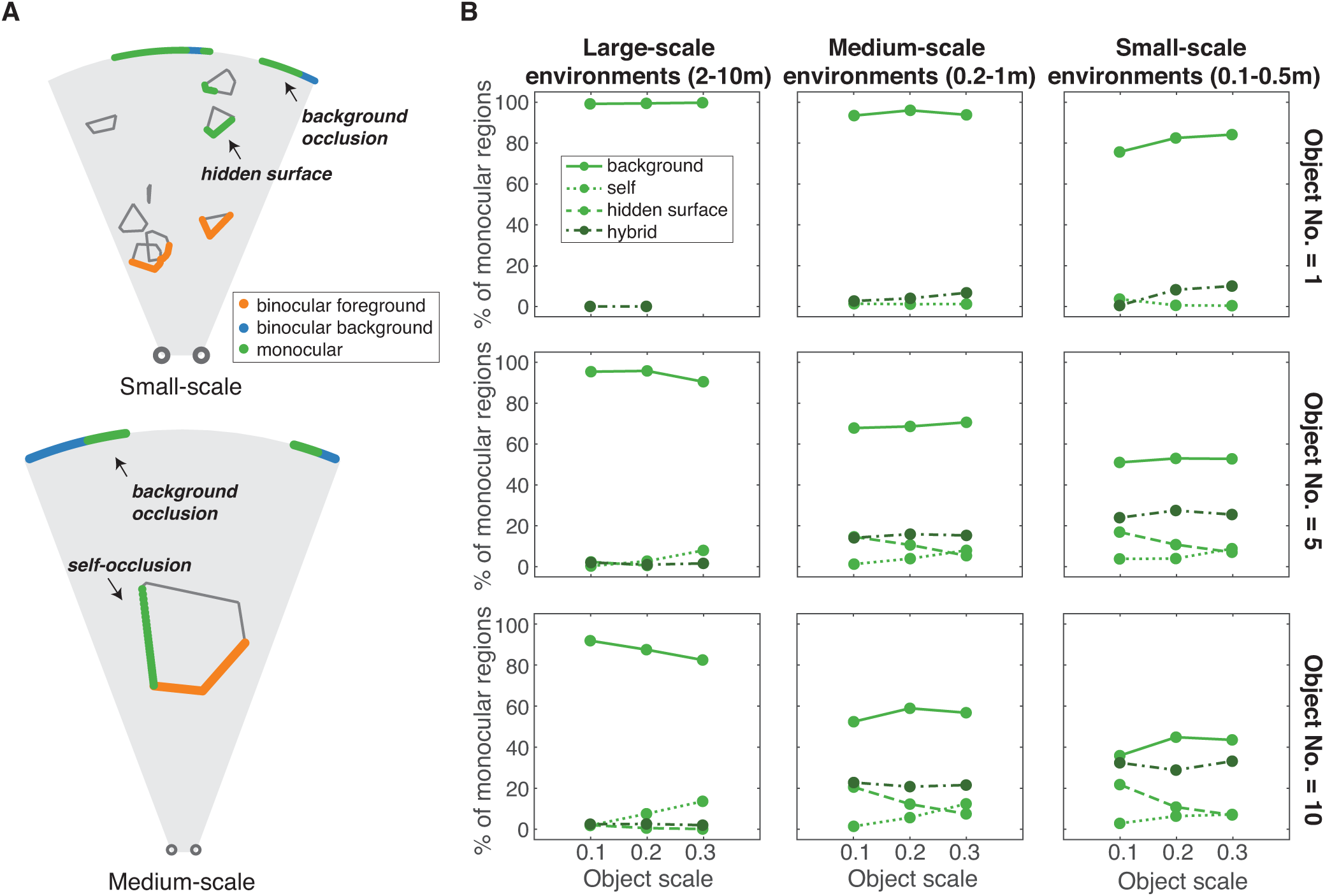
Geometric simulation. **A.** Example diagrams of two simulated environments are shown in a top-down view. The small circles at the bottom illustrate the location of a virtual observer’s eyes (interpupillary distance of 6.5 cm), and the shaded area represents the visible environment. Object size was first specified as the diameter of a circle bounding a polygon, as a proportion of the maximum distance in the current environment (0.1 in upper, 0.3 in lower panel). Random vertices were then placed along each circle, so the objects’ final sizes ranged depending on the vertex locations. The number of vertices was randomly selected to be between 2 and 8, resulting in polygons with between 1 and 7 sides. A minimum object distance was set as 20% of the maximum distance for each environment. The number of objects for each simulation was varied as well. **B.** We ran 200 simulations per parameter combination and determined monocular pixels by ray tracing over the central 40□ field of view at a resolution of 60 rays per degree. We identified all unique monocular regions and categorized each region as being a background occlusion, self-occlusion, hidden surface occlusion, or hybrid occlusion. A hybrid occlusion was one that contained less than 80% of any of the three other categories. The effect of environmental scale (maximum distances of 10, 1, 0.5 m) is shown in different columns, and number of objects (1, 5, 10) in different rows. Each panel shows the percentage of monocular regions in each category, as a function of object scale (0.1, 0.2, 0.3). These scales correspond to sizes that range from 1-3m, 0.1-0.3m, and 0.05-0.15m for the large-, medium-, and small-scale environments, respectively. Data points at 0% are not plotted. Across all parameter combinations, 95% confidence intervals, as determined by the binomial distribution, ranged from 6-11% (not plotted). The different line styles correspond to the four different kinds of monocular regions: background occlusion, self-occlusion, hidden surface occlusion, and hybrid occlusion.

In each simulation, we identified all monocular regions occurring at depth edges within the observer’s central visual field (40□), and categorized each monocular region as being a background occlusion, self-occlusion, hidden surface, or hybrid occlusion (see **Figure 2** and **Figure A1**). A hybrid occlusion was defined as a continuous monocular region that contained less than 80% of any of the three other categories. The results indicate that monocular regions are most likely to be background occlusions across diverse environments with different scales, object densities, and object sizes. Each subpanel of **Figure 8B** summarizes the results for a given environment scale (columns) and object density (rows), and the abscissa of each panel depicts variations in object scale.

We start by examining the results from the large-scale environments, which are most similar to the scenes used in the natural scene statistics (left column). Consistent with the results from our analysis of natural stereo-images, the majority of monocular regions in these simulated environments tended to be background occlusions (solid lines). Other types of occlusions were present, but highly infrequent by comparison. These results suggest a strong constraint on the content of monocular regions when objects are relatively far away from the observer. It would be plausible for this constraint to weaken or reverse when objects are nearby. However, the observation that most monocular regions were background occlusions generalized to smaller scale environments in this simulation as well (right and center column). These smaller environments produced more monocular regions overall, with background occlusions still consistently the most prevalent. Not surprisingly, however, the relative proportion of other types of occlusions was higher in the small-scale environments compared to larger-scale environments. For example, for small-scale environments with many objects, background occlusions and hybrid occlusions had relatively similar frequency. Naturally, as the number of objects increased (upper to lower row), the number of self-occlusions, hidden surfaces, and hybrid occlusions also increased. Interestingly, hidden surfaces and hybrid occlusions were often more frequent than a self-occlusion in this simulation.

Lastly, our results are consistent with the simulations presented by Langer & Mannan (2012), in that we observe more monocular regions when the environments are populated with smaller objects (Langer & Mannan, 2012). For example, in the small-scale environments with the many small objects, over 30% of visible points were monocular, whereas in the large-scale environments less than 3% of visible points were monocular. The assumptions made for these simulations clearly do not fully capture the complex structure of natural scenes. Nevertheless, the assumptions allowed us to parametrically explore how different environmental factors may affect the prevalence and properties of monocularly visible surfaces. These simulations suggest that the monocularly visible regions of real-world scenes are much more likely to belong to background surfaces.

## Discussion

The images that are cast on the retinas during natural viewing carry incomplete information about the three-dimensional scene in front of the observer. However, the visual system often exploits statistical regularities in these images to achieve accurate percepts of the environment. Perception of depth edges may be facilitated by assumptions about the configurations that are likely to occur in natural scenes. The current work provides evidence that there are indeed strong statistical regularities near depth edges in natural scenes, and that these regularities have a systematic effect on perceptual stability.

To examine these perceptual effects, we measured human perceptual stability in two experiments using stimuli taken from natural scenes and analyzed the results with mixed linear regression models. Models that included image features (e.g., edge strength, luminance differences) in addition to disparity as predictors for perceptual instability explained 7-8% more variance than a Disparity-Only model. This is a promising result given the relatively small total effect size and the limitations of the current approach. For example, it is likely that there are stimulus factors (e.g. higher-order textural properties) that are useful for predicting instability and are not well captured by the current set of predictors. In addition, in the current perceptual experiment, all stimuli were presented at a fixed distance (relative to the foreground), even though the natural scene patches were sampled from a range of distances. An exploratory analysis suggested that patches derived from farther distances may tend to be more perceptually stable, perhaps because of differences in textural or perspective cues. The interaction between binocular disparity and textural cues at depth edges represents an interesting direction for future work.

Our current set of models explained only around 1/3 of the variance in mean instability ratings, leaving a substantial proportion of the variance unaccounted for. Indeed, repeated presentations of the same stimulus could elicit quite variable instability ratings within observers. With additional training procedures or a two-alternative forced choice design, this stimulus-independent variability in our response measure may be further reduced. Another potential next step might be to employ more complex textural descriptors derived from natural scenes, or to use automatic techniques for finding the most useful image features for the task (Burge & Jaini, 2017; Geisler, Najemnik, & Ing, 2009; Jaini & Burge, 2017). Doing so may reveal additional stimulus factors that are useful for predicting perceptual instability. Finally, image manipulation may also be used causally manipulate perceptual instability, for example, by introducing luminance and contrast differences between adjacent image regions. However, locally manipulating natural images without producing visible artifacts can be challenging.

The current work has focused on local statistics and low-level image features. In addition to these local statistics, the global structure of a scene is likely to play a role in the perceptual stability of depth edges. For example, a previous study that used photographs of real world objects as stimuli (tabletop scenes with boxes) suggested that correct depth ordering judgements were facilitated for self-occlusions, as compared to background occlusions (Wilcox & Lakra, 2007). Such stimuli may produce a strong expectation that the side of the object should be visible. Recent work also suggests that the detection of depth edges in natural images may be facilitated by taking global layout information (e.g., elevation) into account (Ehinger, Graf, Adams, & Elder, 2017). More global contextual cues may thus lead the visual system to interpret a given scene as more likely to contain one type of occlusion over another, but a thorough understanding of local image cues is clearly important. Early, feedforward stages of visual processing rely on localized computations that are likely made more informative by incorporating low-level statistical regularities from the environment. For example, Anderson & Nakayama (1994) proposed that binocular combination relies on both disparity-tuned receptive fields and receptive fields tuned to detect monocular regions at depth edges at the earliest stages of visual processing. In the search for neural correlates of binocular combination, it has been demonstrated that binocular populations in V1 include neurons that are sensitive to interocular differences that do not typically occur for binocular points during natural vision (e.g., phase disparities) (Ohzawa, DeAngelis, & Freeman, 1990; Read & Cumming, 2007). However, the ability of these populations to detect monocular points is unknown. Recent work on the natural statistics of depth edges may provide guidance for investigating the predicted properties of neurons optimized to encode the combination of monocular and binocular features at depth edges (Iyer & Burge, 2018).

Inspired by the proposed mechanisms for biological vision, some computer vision approaches also aim to simultaneously estimate the disparity of binocular points and locations of monocular points (Wang & Zickler, 2019; Weng, Ahuja, & Huang, 1988). In the field of computer vision, monocular regions have been of interest for their role in facilitating detection of object boundaries when stereo image pairs are available. Recent work in this area has begun to incorporate assumptions about typical scene geometry into this procedure (Wang & Zickler, 2019). However, they have only considered variations of background occlusions, rather than detecting self-occlusions or hidden objects. Based on the primary results presented in the current study, it seems probable that this approach from computer vision would work well on natural stereo images, given that background occlusions are the most likely type of occlusion. Nonetheless, the results of our simulation suggest that hidden object scenarios may also contain useful cues to depth edges, particularly in cluttered scenes.

Looking forward, a systematic exploration of how visual context across multiple scales impacts binocular combination may improve the ability to predict when natural stimuli are most easily fused. In addition, a complete understanding of binocular fusion during natural vision should take into account complex local depth relationships, which can include nested monocular regions associated with multiple half-occluded surfaces (see Assee & Qian, 2007). This understanding would be facilitated by additional datasets from a wider variety of natural scene types (e.g., indoors, outdoors, natural, man-made, etc.). Finally, the temporal dynamics of natural vision, caused by motion in the scene as well as observer motion, likely have a strong modulatory effect on both visual statistics and perceptual stability. As we learn more about the combined features of depth and other cues during natural binocular vision, we can expect that additional sources of information may be discovered that facilitate stable percepts.

## Acknowledgements

The authors would like to thank Steve Cholewiak for equipment assistance. EAC and ZB were supported by Google. JB and DNW were supported by NIH grant R01-EY028571 from the National Eye Institute and the Office of Behavioral and Social Sciences Research. JB was also supported by startup funds from the University of Pennsylvania.

## Appendix

**Figure A1.**
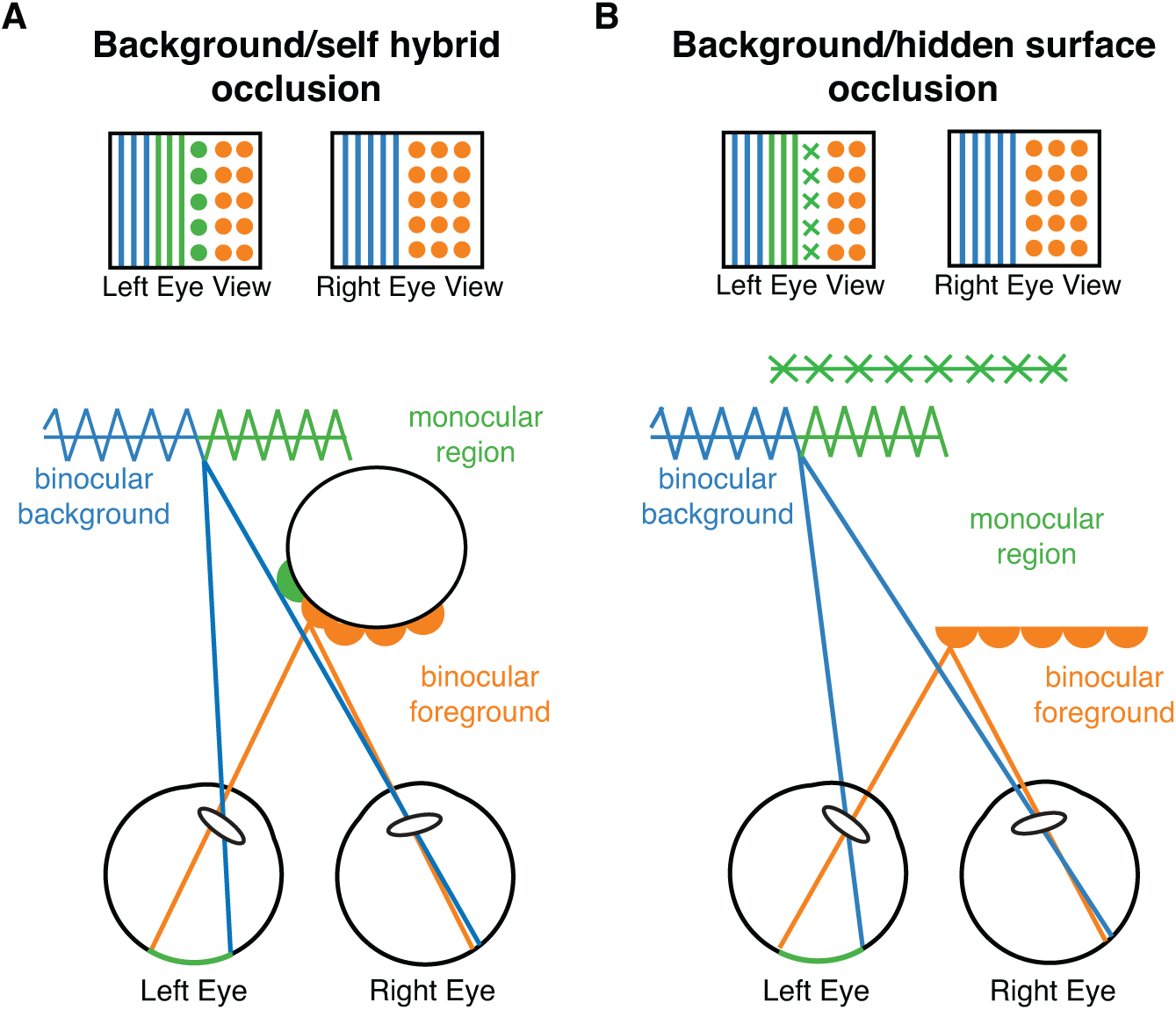
Additional diagrams illustrating cases in which the monocular region has multiple visible surfaces (hybrid occlusions). **A.** In a background/self hybrid occlusion, the left eye sees both a portion of the foreground and background that are hidden from the right eye. **B.** In a background/hidden surface hybrid occlusion, the background is interrupted by a hidden surface, which is also monocularly visible.

